# The human cytomegalovirus ER resident glycoprotein UL148 activates the unfolded protein response

**DOI:** 10.1101/328955

**Authors:** Mohammed N.A. Siddiquey, Hongbo Zhang, Christopher C. Nguyen, Anthony J. Domma, Jeremy P. Kamil

## Abstract

Eukaryotic cells are equipped with three sensors that respond to the accumulation of misfolded proteins within the lumen of the endoplasmic reticulum (ER) by activating the unfolded protein response (UPR), which functions to resolve proteotoxic stresses involving the secretory pathway. Here, we identify UL148, a viral ER resident glycoprotein from human cytomegalovirus (HCMV), as an inducer of the UPR. Metabolic labeling results indicate that global mRNA translation is markedly decreased when UL148 expression is induced in uninfected cells. Further, we find evidence suggesting that ectopic expression of UL148 is sufficient to activate at least two UPR sensors: the inositol requiring enzyme-1 (IRE1), as indicated by splicing of *Xbp1* mRNA, and the PKR-like ER kinase (PERK), as indicated by phosphorylation of eIF2*α* and accumulation of ATF4 protein. During wild-type HCMV infection, *Xbp-1* splicing, eIF2*α* phosphorylation and ATF4 accumulation neatly accompanied the onset of UL148 expression. However, the appearance of these UPR indicators was either markedly delayed or absent during *UL148*-null infections. siRNA depletion of PERK dampened the extent of eIF2*α* phosphorylation and ATF4 induction observed during wild-type infection, implicating PERK as opposed to other eIF2*α* kinases. A virus disrupted for *UL148* showed statistically significant 2- to 4-fold decreases during infection in the levels of transcripts canonically regulated by PERK/ATF4 and by the ATF6 pathway.Taken together, our results argue that UL148 is sufficient to activate the UPR when expressed ectopically and that UL148 is an important cause of UPR activation in the context of the HCMV infected cell.

**IMPORTANCE:** The unfolded protein response (UPR) is an ancient cellular response to ER stress of broad importance to viruses. Certain consequences of the UPR, including mRNA degradation and translational shut-off, would presumably be disadvantageous to viruses, while other attributes of the UPR, such as ER expansion and upregulation of protein folding chaperones, might enhance viral replication. Although HCMV is estimated to express at least 200 distinct viral proteins, we show that the HCMV ER resident glycoprotein UL148 contributes substantially to the UPR during infection, and moreover is sufficient to activate the UPR in non-infected cells. Experimental activation of the UPR in mammalian cells is difficult to achieve without the use of toxins. Therefore, UL148 may provide a new tool to investigate fundamental aspects of the UPR. Furthermore, our findings may have implications for understanding the mechanisms underlying the effects of UL148 on HCMV cell tropism and evasion of cell mediated immunity.

## INTRODUCTION

The endoplasmic reticulum (ER) is a fundamental eukaryotic organelle comprised of a tubulovesicular network of membranes that extends throughout the cytosol [reviewed in (1, 2)]. The organelle carries out multifarious processes vital to cellular and organismal health. For instance, the ER plays key roles in the regulation of intracellular calcium levels, and provides the site for steroid and lipid synthesis, loading of peptides onto MHC complexes (3), and synthesis and processing of proteins and protein complexes destined for secretion. Therefore, it is no surprise that the ER is exploited by a diverse array of viruses during their replication. For instance, polyomaviruses exploit the ER for entry (4, 5), whereas flaviviruses (6) and caliciviruses (7) remodel it to creates sites for replication. Large enveloped dsDNA viruses, such as those in the Herpesviridae, require the ER for the expression and processing of extraordinarily large amounts of viral glycoproteins needed for the assembly of progeny virions.

In order to utilize the ER to support their replication, however, viruses have had to develop mechanisms to contend with the unfolded protein response (UPR), an ancient stress response that serves to maintain ER function and cell viability when misfolded proteins accumulate within the secretory pathway [reviewed in (8, 9)]. The UPR is initiated by three different ER-based signaling molecules: inositol-requiring enzyme-1 (IRE1), the PKR-like ER kinase (PERK), and the cyclic AMP-dependent transcription factor 6 *α* (ATF6). Misfolded proteins are thought to displace the ER chaperone BiP (Grp78) from the luminal domains of IRE1, PERK and ATF6, which causes their activation. During the UPR, mRNA translation is attenuated, and transcripts associated with rough ER ribosomes are degraded, and a number of genes are transcriptionally upregulated, resulting in increased expression of ER protein folding chaperones, ER-associated degradation (ERAD) proteins, as well as various factors that can expand the size and secretory capacity of the ER. Hence, certain consequences of the UPR, particularly translational attenuation, would be expected be deleterious to viruses, while others, such as ER expansion, could enhance the capacity of the infected host cell to produce progeny virions.

Cytomegaloviruses have been found to activate the UPR while subverting certain aspects of it (10, 11). Interestingly, the viral nuclear egress complex component m50 of murine cytomegalovirus (MCMV) degrades IRE1, and the human cytomegalovirus (HCMV) homolog UL50 apparently shares this activity (12). Cytoplasmic splicing of *Xbp-1* mRNA is mediated by IRE1 nuclease activity upon UPR activation. This splicing event is required for translation of the transcription factor XBP1s, which upregulates ERAD factors and ER chaperones, among other target genes (13). In addition, IRE1 degrades mRNAs undergoing translation at the rough ER (14). Therefore, IRE1 downregulation may help to maintain viral glycoprotein expression in the face of UPR activation. Despite this function of UL50, Isler et. al. found evidence that IRE1 is activated during HCMV infection (10). In addition to IRE1, PERK is activated during HCMV and MCMV infection (10, 11), and the PERK / ATF4 axis appears to be required for efficient viral replication, as defects in viral upregulation of lipid synthesis are observed in cells lacking PERK (15).

Interestingly, the viral proteins or processes that activate PERK and IRE1 in the context of HCMV infection have not been clearly identified. We recently reported that UL148 interacts with SEL1L, a component of the cellular ERAD machinery that plays crucial roles in the disposal of misfolded proteins from the ER (16). Having observed very poor expression for any glycoprotein ectopically co-expressed with UL148 in uninfected cells (not shown), we hypothesized that UL148 might trigger the UPR. Here, we show that ectopically expressed UL148 is not only sufficient to activate the PERK and IRE1 arms of the UPR, but also strongly contributes to the activation of PERK and IRE1 during HCMV infection.

## RESULTS

### Ectopic expression of UL148 attenuates translation

As a first step to formally investigate whether UL148 might contribute to ER stress that would trigger the unfolded protein response (UPR), we asked whether ectopic expression of UL148 in uninfected cells would dampen protein synthesis, since translational shutdown is a hallmark of stress responses, including the UPR. To address this question, we employed a “tet-on” lentiviral vector system that would allow us to inducibly express UL148 or its homolog from rhesus cytomegalovirus, Rh159 (17, 18), each harboring a C-terminal influenza A hemagluttinin (HA) epitope tag. Rh159 was used to control for any nonspecific effects of overexpression of an ER resident glycoprotein. We chose Rh159 as a control for the following reasons. Firstly, like UL148, Rh159 is predicted to be type I transmembrane protein with a very short cytoplasmic tail. Secondly, although Rh159 shares 30% amino acid identity with UL148, these two proteins appear to carry out different functions (18-20). Thirdly, UL148 and Rh159 appeared to express at similar levels during ectopic expression (see below).

Having isolated stably transduced ARPE-19 cell populations, we confirmed that that anti-HA immunoreactive polypeptides of the expected size for UL148 (i148^HA^) or Rh159 (i159^HA^) were induced upon treatment with 100 ng/mL doxycycline (dox) (Fig. 1A). Furthermore, expression of neither protein caused any overt reduction in cell viability or number, as measured by trypan blue exclusion following 24 h of dox induction (Fig. 1B, 1C). We therefore concluded that the i148^HA^ and i159^HA^ ARPE-19 cells were suitable to address whether UL148 might affect rates of mRNA translation in metabolic labeling studies. For these experiments, i148^HA^ and i159^HA^ cells were induced (or mock induced) for transgene expression for 24 h, and then incubated in the presence of ^35^S-labeled methionine and cysteine for 30 min. In parallel, labeling was also carried out using i159^HA^ cells that were incubated in the presence of either thapsigargin (Tg) or carrier-alone, so as to provide positive and negative controls, respectively, for UPR induction.

**FIGURE 1.**
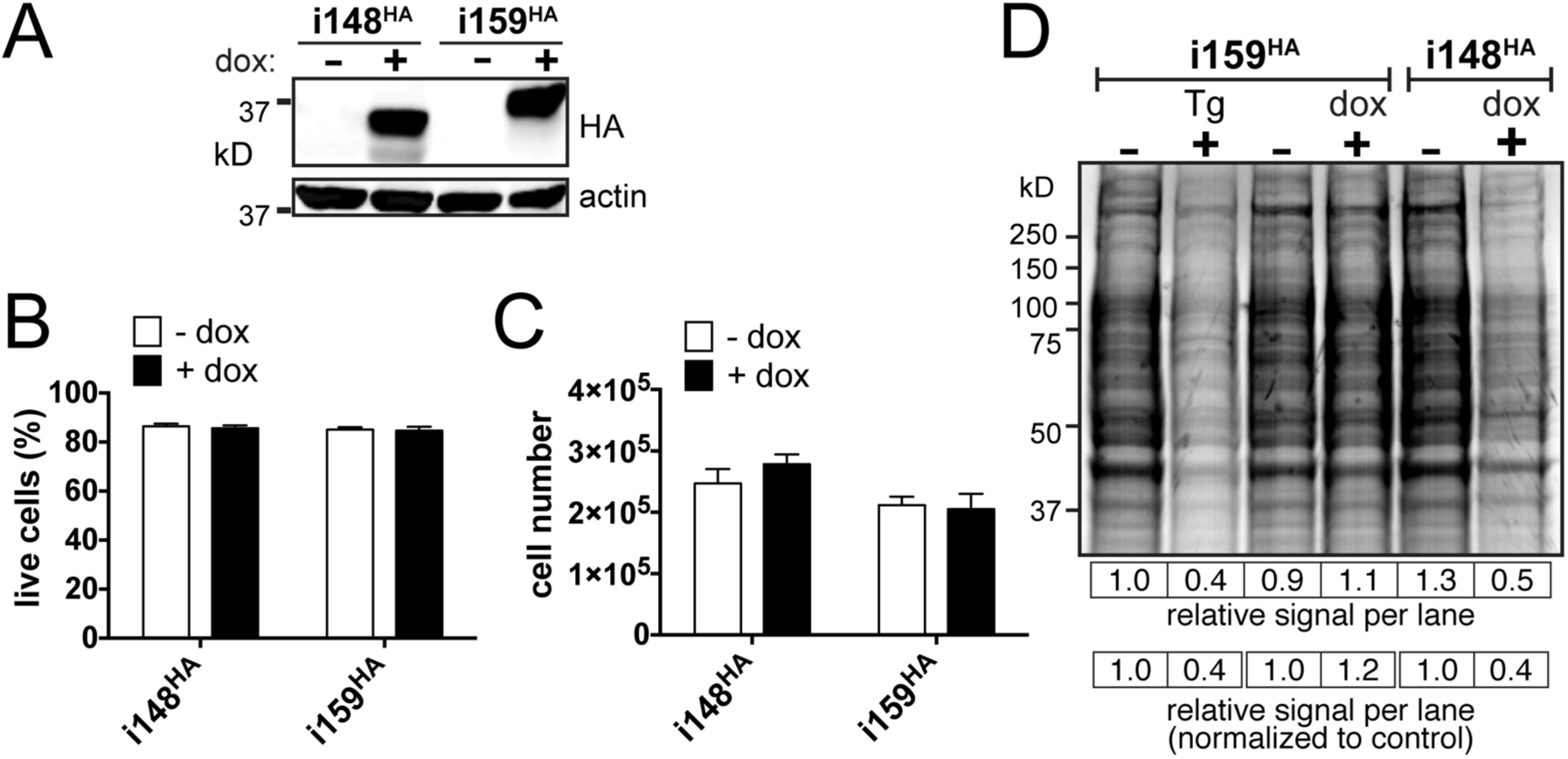
Ectopic expression of UL148 attenuates translation. **(A)** Validation of expression system: i148^HA^ and i159^HA^ ARPE-19 cells were treated with either 100 ng/mL doxycycline (dox) or carrier-alone (water) and subjected to western blot using anti-HA antibodies; beta-actin (actin) was detected as a loading control. **(B)** and **(C)** UL148 expression does not have overtly toxic effects: i148^HA^ and i159^HA^ ARPE-19 cells were dox- or mock induced for 24 h. Viable cells from triplicate treatments were scored using trypan blue exclusion and total cell number on a hematocytometer. **(D)** i148^HA^ and i159^HA^ cells were either dox induced or mock treated (0.1% water) for 24 h, and in parallel, additional wells of i159^HA^ cells were treated for 2 h with either 2 µM thapsigargin (Tg) or 0.1% DMSO carrier-alone. Cells were then pulsed with ^35^S methionine +cysteine for 30 min, and protein lysates were resolved by SDS-PAGE and imaged by autoradiography. Quantitation of the total signal intensity per lane is shown relative to that of the (-) Tg lane (leftmost) directly below the gel image and for each treatment lane relative to its paired negative control.

We found that expression of UL148 but not Rh159 caused a substantial, ∼50% decrease in protein synthesis compared to the carrier-alone (water) treated control, as measured by phosphor-image analysis (Fig. 1D). Strikingly, the attenuation of translation observed during UL148 expression was similar in magnitude to that seen during Tg treatment (Fig. 1D). These effects did not appear to be caused by the inducing agent, since dox treatment of i159^HA^ cells failed to cause any reduction in ^35^S incorporation. From these results, we concluded that expression of UL148 attenuates translation. Since UL148 is an ER resident glycoprotein, with a predicted type I transmembrane topology that places most of the polypeptide in the ER lumen (19), it seemed plausible that the effects of UL148 on global rates of mRNA translation might be indicative of the UPR. We thus sought to address the hypothesis that UL148 activates the UPR.

### UL148 leads to PERK-dependent phosphorylation of eIF2*α* and accumulation of ATF4

Translational attenuation during the UPR is mediated by the PKR-like ER kinase, PERK, which phosphorylates Ser51 of the *α*-subunit of the ternary eIF2 complex (21). The guanine nucleotide exchange factor eIF2B binds to the phosphorylated eIF2 complex with increased affinity, and fails to exchange bound GDP for GTP (22). Since GDP/GTP exchange is necessary for eIF2 to participate in a new round of translational initiation, and because eIF2*α* is present in cells at a considerable molar excess relative to eIF2B, global protein synthesis halts in response to even modest levels of phosphorylated eIF2*α* [reviewed in (23)]. Meanwhile, eIF2*α* phosphorylation leads to enhanced translation of certain mRNAs, such as that encoding ATF4, which harbor µORFs in their 5’UTRs that inhibit their translation under non-stressed conditions (24). Although there are four different kinases that have been identified to phosphorylate eIF2*α* at Ser51, two observations imply that the translational attenuation we observed during UL148 expression was due to activation of PERK: (i) UL148 localizes to the ER (19), and (ii) interacts with the ERAD machinery (16). Therefore, we next monitored levels of PERK, eIF2*α* phosphorylation, and ATF4 following dox induction of either UL148 or Rh159 in ARPE-19 cells.

We observed that UL148 and Rh159 proteins accumulated to readily detectable levels by 8 h post induction with dox, although faint expression was detected at 4 h post induction (Fig. 2). By 24 h post induction, the i148^HA^ cells showed robust levels of ATF4 protein, albeit not as high as those seen during Tg treatment, which was included as a positive control for PERK activation. Increased levels of phospho-eIF2*α* were detected from 24 h to 48 h following induction of UL148, but not during induction of Rh159. Moreover, PERK protein levels appeared to be upregulated at 24 h post induction in i148^HA^ cells, but not in i159^HA^ cells. Decreased mobility of the anti-PERK immunoreactive band, which likely indicates PERK autophosphorylation upon UPR activation, was readily observed in the Tg condition, but not following induction of either UL148 or Rh159 (Fig. 2), which may indicate that PERK is less synchronously activated following dox-induction of UL148 than by the comparatively shorter (4 h) Tg treatment. Although we could not exclude the possibility that UL148 might cause these effects via activation of different eIF2*α* kinase, the simplest interpretation of these results is that expression of UL148 activates PERK.

**FIGURE 2.**
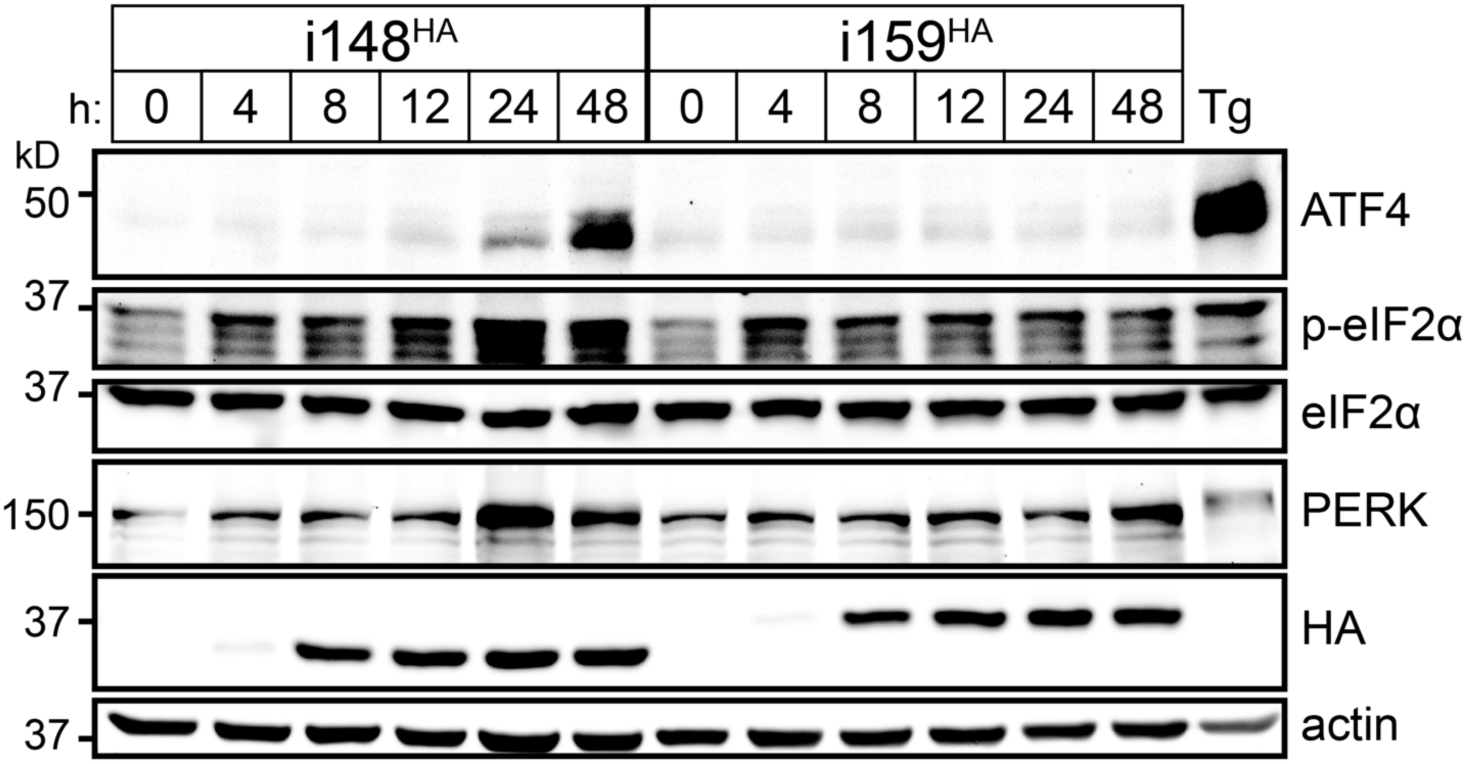
UL148 expression causes phosphorylation of eIF2*α* and accumulation of ATF4, suggesting activation of PERK. i148^HA^ and i159^HA^ ARPE-19 cells were induced for expression of UL148 and Rh159, respectively, using 100 ng/mL doxycycline (dox) for the indicated times. A 4 h treatment with thapsigargin (Tg) (0.4 µM) was included as positive control. For each sample, a volume of lysate equivalent to 17.5 µg of detergent soluble protein was analyzed by western blot for levels of the indicated proteins.

### UL148 is sufficient to induce splicing of *Xbp-1* mRNA

To determine whether UL148 activates IRE1, we transfected human embryonic kidney (HEK)-293T cells with plasmids that drive expression of UL148 or Rh159 carrying C-terminal HA-tags. We also examined the effects of a 2 h treatment with 1 mM dithiothreitol (DTT), as a positive control treatment known to activate IRE1. At 48 h post transfection, we harvested cells for isolation of total RNA and for protein lysates to monitor transgene expression. As a read-out for IRE1 activity, we used reverse-transcriptase PCR (RT-PCR) to detect the removal of 26 nucleotides (nt) from the *Xbp-1* mRNA. This unorthodox splicing event is catalyzed in the cytosol by IRE1; its detection is widely used as an indicator of the UPR in general, and of IRE1 nuclease activity in particular (25-27). Although we also tried this assay using our dox-inducible ARPE-19 cells (not shown), we found that transient transfection of HEK-293 cells gave the most readily interpretable results (Fig. 3).

**FIGURE 3.**
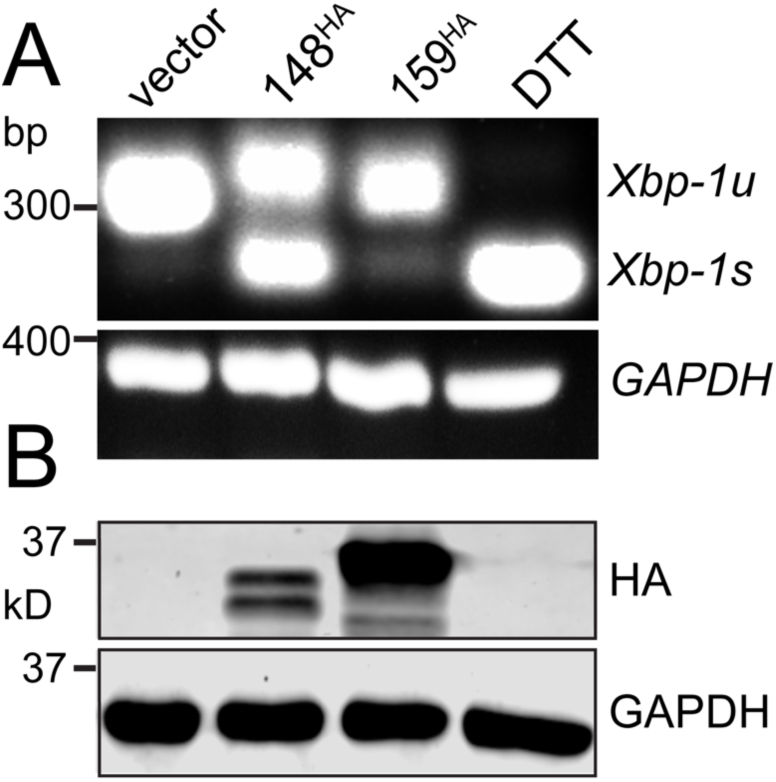
UL148 induces splicing of *Xbp-1* mRNA. HEK-293T cells were transfected with 1 µg of plasmid DNA for the indicated expression constructs or with empty vector as a negative control. **(A)** 48 h post transfection, total RNA was isolated and used in an RT-PCR assay to detect removal of the 26 nt intron from *Xbp-1* mRNA. *Xbp-1s*: spliced message, *Xbp-1u*: unspliced message. A 2 h treatment with dithiothreitol (DTT, 1 mM) was used as positive control. **(B)** UL148 and Rh159 were detected using anti-HA western blot. In both panels, *gapdh* mRNA or GAPDH protein was detected as a loading control.

As expected, the 2 h DTT treatment caused the 26-nt intron to be spliced from nearly all the *Xbp-1* mRNA detected in our assay (Fig. 3A). Cells transfected with either the Rh159 expression plasmid or empty vector failed to show notable levels of *Xbp-1* splicing. In contrast, removal of the 26-bp was readily detected from cells expressing UL148, with approximately equal levels of RT-PCR products for spliced and unspliced *Xbp-1* (Fig. 3A). Furthermore, the expression of anti-HA immunoreactive bands of the expected sizes for Rh159 and UL148 was confirmed by western blot (Fig. 3B). From these results, we concluded that ectopic expression of UL148 but not Rh159 is sufficient to induce splicing of the 26-nt intron from *Xbp-1* mRNA. Given that IRE1 is required for this splicing event (25-28), our results argue that UL148 expression is sufficient to activate IRE1.

### UL148 activates IRE1 during HCMV infection

Since UL148 was apparently capable of activating the UPR when ectopically expressed, we wondered whether UL148 might contribute to the UPR activation in the context of HCMV infection. Therefore, we conducted a time-course experiment comparing IRE1 catalyzed splicing of *Xbp-1* in fibroblasts infected at MOI 1 with either wild-type (WT) HCMV strain TB40/E (TB_WT) or a *UL148*-null mutant, TB_148_STOP_ (16). Remarkably, while WT infected cells showed increasing levels of spliced *Xbp-1* (*Xbp-1s*) as infection progressed, *UL148*-null virus infected cells, showed only very low levels of spliced *Xbp-1* that did not increase over time (Fig. 4). It is also notable that during WT infection the proportion of spliced to unspliced *Xbp-1* message increased from 24 hpi to 144 hpi. These effects correlate nicely with appearance of detectable levels of UL148 during infection and with increases in its levels that occur as infection progresses (Fig. 5). Further, we observed comparable levels of *IE2* mRNA by semi-quantitative RT-PCR, indicating that infection with the two viruses occurred at similar levels (Fig. 4). We interpreted these results to suggest that UL148 contributes to activation of IRE1 and concomitant cytoplasmic splicing of *Xbp-1* mRNA during HCMV infection. Since *Xbp-1* splicing is an important hallmark of UPR activation, these results also suggest that UL148 is a considerable source of ER stress in the context of HCMV infected cells.

**FIGURE 4.**
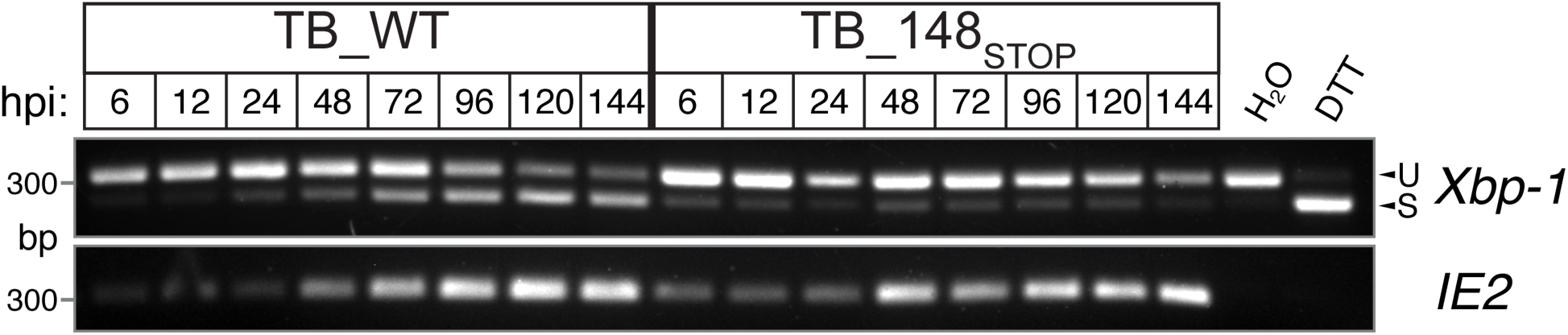
UL148 activates IRE1 during HCMV infection. Fibroblasts were infected MOI of 1 TCID_50_ per cell with wild-type HCMV strain TB40/E, (TB_WT) or a *UL148-*null mutant virus, (TB _148_STOP_). At the indicated times post infection, total RNA was harvested and subjected to RT-PCR to detect IRE1 catalyzed removal of the 26 bp intron from the *Xbp-1* mRNA. Semi-quantitative RT-PCR of *IE2* (*UL122*) is shown as a control to indicate HCMV gene expression. U: unspliced, S: spliced. DTT: dithiothreitol, a positive control.

**FIGURE 5.**
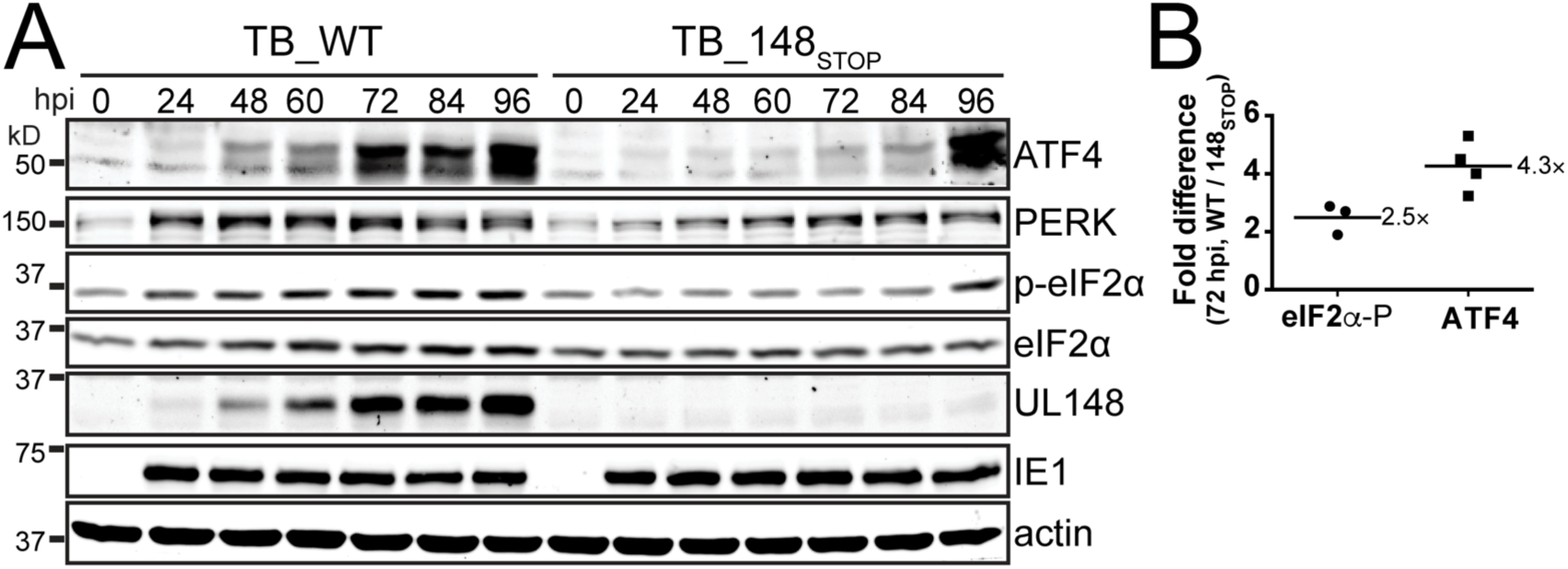
UL148 is required for increases in eIF2α phosphorylation and ATF4 protein expression during HCMV infection. **(A)** Fibroblasts were infected MOI of 1 TCID50 per cell with TB_WT or TB_148_STOP_ viruses for the indicated times and subsequently assayed by western blot (25 µg of protein per lane) for levels of ATF4, PERK, p-eIF2α, eIF2α, UL148, IE1, and beta-actin. **(B)** Fold-difference in phospho-eIF2α (p-eIF2α) and ATF4 protein signals at 72 hpi between TB_WT or TB_148_STOP_. Fluorescence signal from secondary antibodies at the 72 hpi time points was quantified from western blots comparing TB_WT and TB_148_STOP_ infections, as in (**A**), using three independent biological replicates for p-eIF2α and four biological replicates for ATF4.

### UL148 contributes to PERK-dependent increases in phosphorylated eIF2*α* and ATF4 during HCMV infection

To evaluate whether UL148 contributes to PERK activation during infection, we monitored levels of PERK, phospho-eIF2*α*, and ATF4 following MOI 1 infection of fibroblasts with WT or *UL148*-null virus. We found striking differences between the WT and *UL148*-disrupted infection contexts in each of these parameters, which together suggest a role for UL148 in activation of PERK. In WT infected cells, ATF4 levels showed an obvious increase at 48 h post infection (hpi) and reached near maximal expression at 72 hpi, with the highest levels detected at 96 hpi (Fig. 5A). The changes in ATF4 expression during WT infection coincided with increased phosphorylation of eIF2*α* and higher levels of PERK, as expected (10, 24). The kinetics of ATF4 expression tightly correlated, to a remarkable degree, with those seen for UL148 (Fig. 5A).

In cells infected with *UL148*-null virus, ATF4 was weakly expressed at most of the time points monitored, although faint increases were seen at 72 hpi and 84 hpi. At 96 hpi, however, a strong burst of ATF4 expression was detected, which was accompanied by an increase in phosphorylated eIF2*α*. This observation suggests that UL148-independent activation of one or more eIF2*α* kinases occurs at very late times during infection. Importantly, levels of the viral IE1 (IE1-72) protein were similar across all time points for both viruses, indicating that infection occurred efficiently in both WT and *UL148*-null settings (Fig. 5A), as would be expected since *UL148*-null mutants replicate indistinguishably from WT in fibroblasts (19). Because the UL148-indepenent rise in levels of phospho-eIF2*α* and ATF4 occurred between 84 and 96 hpi, we reasoned that the 72 hpi time point would best allow us to isolate the effect of UL148 on these indicators of PERK activation. By measuring the fluorescence signal from secondary antibodies from multiple biological replicates, we were able to estimate that at 72 hpi WT infected cells contain 2.5-fold higher levels of phospho-eIF2*α*, and 4.3-fold higher levels of ATF4 relative to UL148_STOP_ infections (Fig. 5B).

In order to more specifically address whether PERK is required for UL148 to cause phosphorylation of eIF2*α* and accumulation of ATF4, we used siRNA to silence PERK expression prior to infection, and then monitored for phosphorylation of eIF2*α* and expression of ATF4 from 24 to 72 hpi. Levels of PERK were substantially reduced but not completely eliminated by the PERK-targeted siRNA treatment, as compared to the non-targeting control siRNA (NTC) (Fig. 6). In PERK-silenced cells during WT infection, phosphorylation of eIF2*α* was attenuated at both 48 hpi and 72 hpi and a substantial decrease in ATF4 was seen at 72 hpi (Fig. 6). During *UL148*-null infections, however, PERK knockdown led to only minimal effects on phosphorylation of eIF2*α*, and virtually imperceptible effects on ATF4 (Fig. 6), which may well reflect reduced levels of ER stress in the absence of UL148.

**FIGURE 6.**
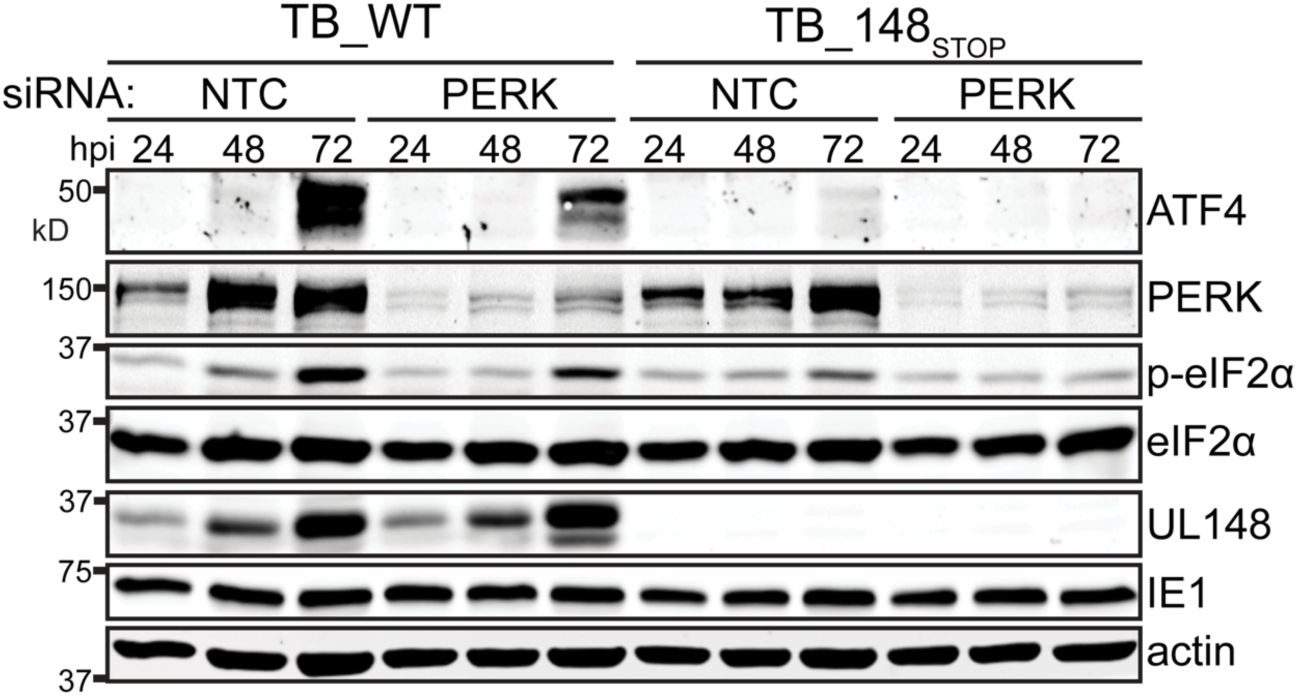
PERK contributes to the effects of UL148 on eIF2α phosphorylation and ATF4 protein levels. Fibroblasts were reverse-transfected using an siRNA SMARTpool targeting PERK or a non-targeting control siRNA pool (NTC). 24 h later, the cells were infected at an MOI of1 TCID50 per cell with either TB_WT or TB_148_STOP_ viruses for the indicated times. Cell lysates were assayed by immunoblot for expression of the indicated proteins.

Quantification of fluorescent secondary antibody signals suggested that in the case of WT virus at 72 hpi, PERK knockdown led to a 40% decrease in the levels of phospho-eIF2*α* and ATF4. Because the siRNA knockdown of PERK was incomplete, these results seem likely underestimate the degree to which UL148 depends on PERK to cause phosphorylation of eIF2*α* and to increase ATF4 expression. Overall, we interpreted these findings to argue that UL148 activates PERK during HCMV infection.

### UL148 contributes to differences in mRNA levels for UPR target genes

A major function of the UPR is to cause changes in cellular gene expression. Since ATF4 and XBP1s are transcription factors that contribute to UPR mediated changes in gene expression (13, 24, 29), we next wished to determine whether UL148 contributes to effects of HCMV on mRNA levels for cellular genes that are known to be regulated by the UPR. Further, because we were unable to obtain antibody sensitive enough to test whether activation of ATF6 was influenced by UL148 (not shown), and because it has been reported that HCMV infection does not lead to ATF6 activation, but that genes regulated by ATF6 are nonetheless upregulated (10, 30), we also sought to address whether UL148 might contribute to upregulation of ATF6 target genes. Therefore, we isolated total RNA from WT and *UL148*-null infected fibroblasts at 72 hpi and used RT-qPCR to measure mRNA levels for representative UPR target genes, including ATF6 target genes in addition to those regulated by ATF4 (PERK) and XBP1s (IRE1).

With regard to the PERK pathway, our results show that relative to *UL148*-null virus infected cells, WT virus infected cells exhibited nearly 4-fold higher mRNA levels for the ATF4 target gene CHOP, and roughly 2-fold higher mRNA levels for another ATF4 target, GADD34 (Fig. 7). These differences were found to be statistically significant (*P*<0.05). Despite the UL148-dependent effects we observed on *Xbp-1* splicing (Figs. 3-4), the levels of mRNAs from XBP1s target genes did not appreciably differ between WT and *UL148*-null infections (Fig. 7). This result is consistent with a previous report that failed to find an effect of HCMV-induced *Xbp-1* splicing on mRNA levels for the XBP1s target gene *EDEM1* (10). Intriguingly, we did find significant differences for a number of ATF6 target genes that were upregulated in WT relative to *UL148*-null infections, including *BiP, SEL1L, HERPUD1*, and *HYOU1*, all of which showed approximately 2-fold higher expression during WT infection. Although *PDIA4* was upregulated by over two-fold in cells infected with WT virus compared to those infected with *UL148*-null virus, this difference was not found reach statistical significance. From these results, we concluded that UL148 contributes during HCMV infection to upregulation of UPR target genes related to the PERK and ATF6 arms of the UPR.

**FIGURE 7.**
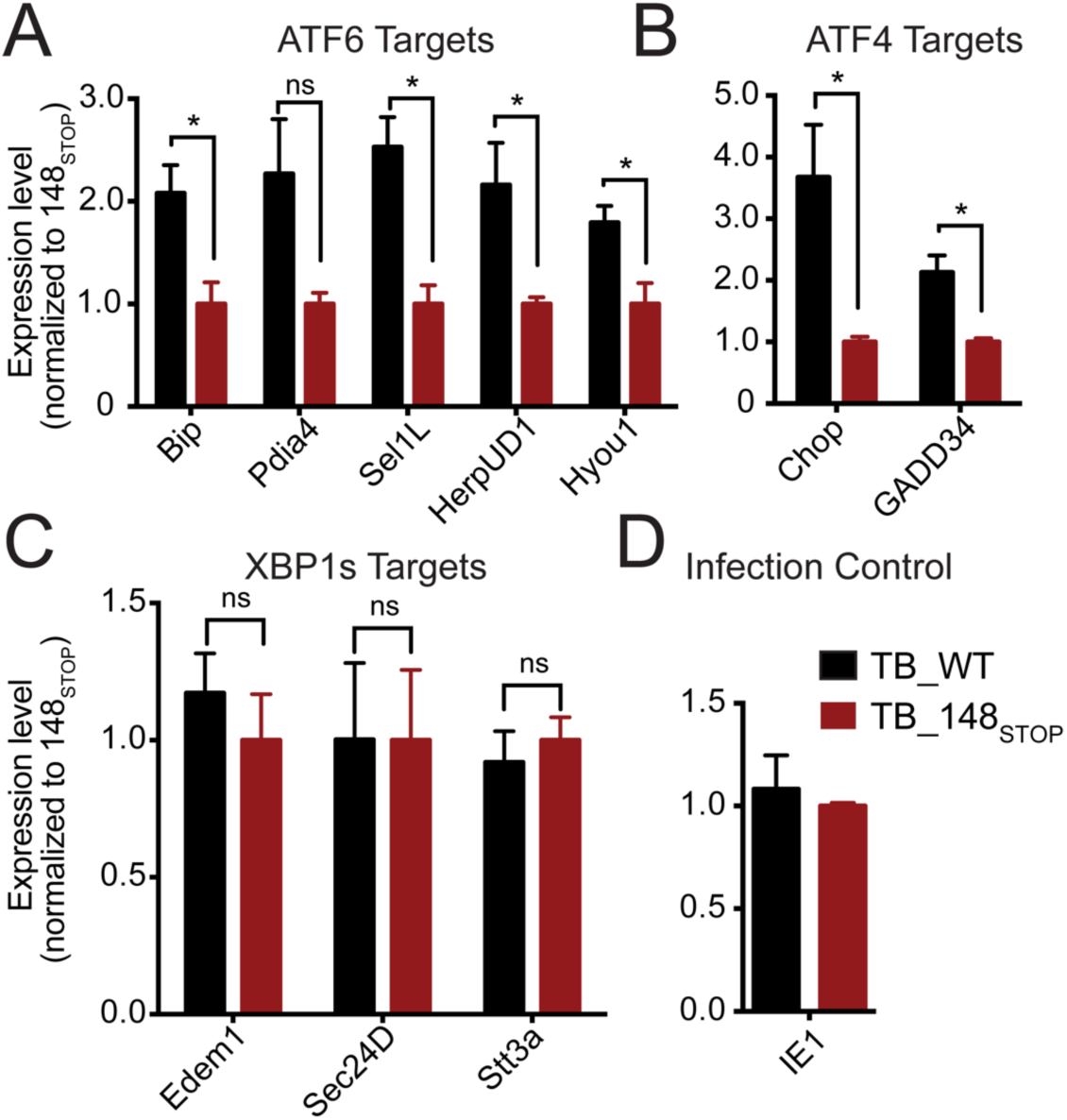
Analysis of mRNA levels for UPR target genes during WT and *UL148*-null infection. Fibroblasts were infected at an MOI of 1 TCID_50_ per cell with TB_WT or TB_148_STOP_ viruses. At 72 h post infection, mRNA levels for the indicated genes were analyzed by quantitative RT-PCR. Asterisks indicate differences found to be statistically significant in a two-tailed T-test (P<0.05).

## DISCUSSION

HCMV is estimated to encode 164-192 distinct genes (31, 32), with a more recent study arguing for up to 751 protein coding ORFs (33). Thus, the degree to which our results suggest that UL148 alone contributes to UPR induction during HCMV infection is remarkable. The original work demonstrating that HCMV activates the UPR conducted their studies using the laboratory-adapted virus strain Towne (10), which unlike another widely studied laboratory strain, AD169, retains the capacity to express UL148 (34, 35). Accordingly, the kinetics of ATF4 protein accumulation and phosphorylation of eIF2*α* that we observed for cells infected with wild-type (WT) strain TB40/E (Fig. 5) are highly consistent with those observed in the latter study (10), and the extent to which these indicators of PERK activation were dampened during *UL148*-null infection is striking (Figs. 5-6).

Cells infected with *UL148*-null viruses exhibited reduced levels of eIF2*α* phosphorylation and impaired induction of ATF4 at times prior to 96 hpi (Fig. 5). Although there are three other eIF2*α* kinases, we contend that because UL148 is an ER-resident protein, and also appears to activate IRE1, another sensor of ER stress, the effects on eIF2*α* phosphorylation and ATF4 levels most like occur via PERK. In support of this notion, PERK knockdown diminished the effect of UL148 on ATF4 induction (Fig. 6). Similarly, in studies with MCMV Qian et al. found that knockdown of PERK led to attenuated levels of ATF4 (11). Thus, both human and murine cytomegaloviruses appear to induce phosphorylation of eIF2*α* and ATF4 upregulation via PERK even though MCMV does not encode a UL148 homolog.

Furthermore, we detected 2 to 4-fold higher mRNA levels for two ATF4-regulated regulated genes, CHOP and GADD34, in WT compared to *UL148*-null infected cells (Fig. 7). As opposed to the effects of UL148 on *Xbp-1* splicing (Fig. 4), which were not accompanied by differences in mRNA levels for XBP1s target genes, the effects of UL148 on the PERK-ATF4 axis were accompanied by the expected changes in gene expression (Fig. 6). Nonetheless, our *Xbp-1* splicing results argue that UL148 is sufficient to activate IRE1 (Fig. 3). Moreover, UL148 appears to account for most of the IRE1 activation observed during HCMV infection (Fig. 4). These effects are particularly noteworthy as they presumably occur in the face of viral downregulation of IRE1 by the viral nuclear egress factor UL50 (12).

Although the kinetics of *Xbp-1* splicing we observed during WT infection were similar to those seen by Isler et al. (10), the ratio of spliced-to-unspliced message appeared to be much higher in our results and may reflect differences in UL148 expression between strains Towne and TB40/E. In our hands, the Towne strain appears to express UL148 at much lower levels than TB40/E (H. Zhang and J.P. Kamil, unpublished results). Regardless, *XBP1s* target genes such as EDEM1 (36, 37) were not found to be upregulated in a UL148-dependent manner (Fig. 6), as is fully consistent with the findings of Isler et al. (10), who likewise failed to observe EDEM1 upregulation despite observing *Xbp-1* splicing during infection.

Since XBP1s target genes are apparently refractory to IRE1 activation during HCMV infection, the implications to the virus of IRE1 activation are unclear. However, splicing of *Xbp-1* is not the only function of IRE1. IRE1 also activates the JNK signaling pathway (38) and degrades mRNAs associated with the rough ER (14). Furthermore, IRE1 confers resistance to apoptosis during hepatitis C infection by degrading mIR-125a (39). Albeit that we used *Xbp-1* splicing as a specific readout for activation of IRE1, it seems conceivable that functions of IRE1 unrelated to splicing of *Xbp-1* mRNA may be relevant to phenotypes governed by UL148.

Whether UL148 activates the third UPR sensor, ATF6, remains unresolved. *UL148*-null virus infected cells did show lower mRNA levels for ATF6 target genes compared to WT infected cells (Fig. 7), which may suggest that ATF6 is activated by UL148. Unfortunately, we have not been able to directly evaluate this matter directly, owing to the limited sensitivity in our hands of commercially available ATF6 antibodies (not shown). ATF6 is proteolytically processed by the same proteases that regulate sterol responsive element binding proteins (SREPBs), S1P and S2P (40). Under conditions of ER stress, ATF6 transits from the ER to the Golgi where S1P and S2P release the cytoplasmic domain of ATF6 from its transmembrane anchor, allowing it to transit to the nucleus where it binds to cis-acting ER stress regulatory elements (ERSE) and upregulates genes for ER chaperones, such as BiP (Grp78) (26, 41).

Nonetheless, upregulation of BiP reportedly occurs in an ERSE-independent manner during HCMV infection (30). Although Isler et al. were unable to detect ATF6 cleavage despite finding target genes to be upregulated during HCMV infection (10), the S1P/S2P processed nuclear form of ATF6, like SREBPs, is rapidly degraded in the absence of proteasome inhibitors (40). Hence, it is difficult to exclude the possibility that that low levels of ATF6 activation occur during HCMV infection. Given these circumstances, it may be challenging to address the potential role of UL148 in ATF6 activation.

### Why would HCMV encode a protein that activates the UPR?

It is intriguing to consider why HCMV would encode a viral protein that potently triggers the UPR. *C*ertain consequences of the UPR, such as enhanced ERAD and attenuation of translation, might be expected to be unfavorable for viral replication. For instance, degradation of mRNAs by IRE1 could hamper the expression of viral glycoproteins, and phosphorylation of eIF2*α* by PERK could dampen viral gene expression. However, maintaining translation of viral mRNAs is a *sine qua non* for cytolytic viruses, and the literature resoundingly suggests HCMV is no exception [reviewed in (42)]. In particular, UL38 plays a role in disarming ER stress, as it appears to limit both PERK activation and stress-induced translation of *ATF4* mRNA during infection (43). Nonetheless, PERK is required for efficient HCMV replication, and defects in viral upregulation of lipid synthesis have been observed during infection of PERK depleted cells (15). Meanwhile, activation of ATF6 and IRE1 are required for ER expansion, upregulation of ER chaperones, and for increased synthesis of lipids (25, 41, 44-48), all of which may benefit viral replication.

Given the substantial contribution of UL148 to UPR activation documented here, and the potential for the UPR to both negatively and positively impact viral replication, it is puzzling that *UL148*-null viruses are found to replicate indistinguishably from WT virus in fibroblasts (16, 19). Although we cannot yet exclude whether decreased induction of the UPR contributes to the enhanced growth of *UL148*-null virus in epithelial cells, the influence of UL148 on the expression of alternative gH/gL complexes, particularly gH/gL/gO, (16, 19) seems a more likely explanation. A derivative of the HCMV strain AD169 that was restored both for *UL148* and for expression of the pentameric gH/gL/UL128-131 complex (16) appears to replicate at least as well in epithelial cells as the parental virus lacking *UL148*, while failing to show differences in gH/gL/gO expression in virions (Nguyen C.C., Siddiquey, M.N.A., Li, G. and Kamil J.P., unpublished results). We thus consider it unlikely that expression of UL148 is directly detrimental to productive replication of HCMV in epithelial cells, especially since viral factors such as UL50 (12) and UL38 (43, 49) would be expected to blunt any negative impacts to the virus of UPR induction.

### Implications for mechanisms underlying *UL148* dependent phenotypes

Going forward, it will be crucial to delineate which biological roles and/or phenotypic effects of UL148 require induction of the UPR. We cannot yet dismiss the possibility that UPR induction is incidental to the *bona fide* biological function(s) of UL148, which could be modulation of virion cell tropism (19), or evasion of cell-mediated immune responses (20). In other words, UPR activation may not be required for the effects of UL148 that provide a fitness advantage to the virus. For example, although the HCMV ER resident immune-ëvasin US11 triggers the UPR in uninfected cells, the UPR does not appear to be required for US11-mediated degradation of the MHC I heavy chain (50). On the other hand, certain observations suggest that UPR induction may be inseparable from the role of UL148 in cell tropism. We recently reported that UL148 co-purifies from infected cells with SEL1L, a key component of the cellular machinery for ER associated degradation (ERAD), and have found that UL148 attenuates ERAD of newly synthesized glycoprotein O (gO), which itself appears to be a constitutive substrate for ERAD (16). Therefore, one might hypothesize that UL148 interacts with the ERAD machinery to impede processing of misfolded proteins, which consequently results in UPR activation.

UL148 was recently found to block surface presentation of CD58 (LFA3), a costimulatory ligand that potentiates T-lymphocyte and NK-cell responses (20). Intriguingly, UL148 causes markedly reduced N-glycosylation of CD58 (20), which is exactly the opposite of its effect on gO (16, 19). Rh159, which shares significant sequence homology with UL148 and is involved in retention of a distinct set of costimulatory molecules(18), does not appear to activate the UPR (Figs. 1-3). Although it is unknown whether UL148 requires UPR activation to downregulate CD58, knowledge of the proximal events by which UL148 activates the UPR will likely prove integral to understanding the mechanisms underlying its influence on viral immune evasion and modulation of tropism.

Finally, it is worth pointing out that UL148 may hold promise as a reagent to investigate the UPR itself. Much of our understanding of the mammalian UPR comes from experimental approaches in which toxic chemicals, such as thapsigargin or tunicamycin, are relied upon to synchronously and robustly induce the UPR in cultured cells. A recent report found that such chemicals fail to accurately recapitulate the authentic UPR induced by unfolded proteins within the ER lumen (51). Albeit that the molecular events by which UL148 initiates the UPR remain to be determined, this viral ER resident glycoprotein may represent a fascinating new tool to interrogate how cells adapt to ER stress.

## MATERIALS AND METHODS

### Cells and Virus

Primary Human foreskin fibroblasts (HFF, ATCC #SCRC-1041) were immortalized by transducing lentivirus encoding human telomerase (hTERT) to yield HFFT cells. HEK-293T cells were purchased from Genhunter Corp. (Nashville, TN). The retinal pigment epithelial cell line ARPE-19 was purchased from ATCC (CRL-2302). All cells were cultured in Dulbecco’s Modified Eagle’s Medium (DMEM, Corning # 10013CV) supplemented with 25 μg/mL gentamicin, 10 μg/mL ciprofloxacin-HCl, and either 5% fetal bovine serum (FBS, Sigma-Aldrich #F2442) or 5% newborn calf serum (NCS, Sigma-Aldrich #N4637).

Viruses were reconstituted by electroporation of HCMV bacterial artificial chromosomes (BACs) into HFFTs, as described previously (19, 52), and grown until 100% CPE was observed. Cell-associated virus was released by Dounce-homogenization of pelleted infected cells, clarified of cell debris by centrifugation (1000*g*, 10 min), and combined with the culture supernatants. Cell-associated and cell-free virus containing virus were combined and then ultracentrifuged through a 20% sorbitol cushion (85,000*g*, 1 h, 4˚C). The resulting virus pellet was resuspended in DMEM containing 20% NCS. Viruses for this study were all derived from the bacterial artificial chromosome clone of HCMV strain TB40/E, TB40E-BAC4 (53), which was a generous gift of Christian Sinzger (Ulm, Germany). A *UL148*-null mutant derived from TB40E-BAC4, TB_148_STOP_, has been described elsewhere (16). BACs and plasmid DNAs for transfection were purified from *E. coli* using Nucleobond Xtra Midi kits (Machery-Nagel, Inc.).

### Virus titration

Infectivity of virus stocks and samples were determined by the tissue culture infectious dose 50% (TCID_50_) assay. Briefly, serial dilutions of virus were used to infect multiple wells of a 96-well plate. After 9 days, wells were scored as positive or negative for CPE, and TCID_50_ values were calculated according to the Spearman-Kärber method, as described previously (16).

### Construction of plasmids

UL148 and Rh159 were PCR amplified from plasmids pEF1-UL148HA (19) and pcDNA-Rh159 IRES-GFP (a gift of Klaus Frueh, Oregon Health Sciences University, Beaverton, OR) using primer pairs UL148_reclone_Fw and UL148 reclone Rv, and Rh159 Fw and Rh159_HA_Rv, respectively (Table 1). The PCR product for UL148 was ligated into pcDNA3.1(+) (Invitrogen) using the BamHI and EcoRI sites, while the PCR product for Rh159 was inserted into the EcoRV site using a Gibson Assembly Reaction using NEB HiFi DNA assembly Master Mix (New England Biolabs). Final plasmids were sequence confirmed using T7 and BGH reverse primers. To construct lentiviral vectors for inducible expression of Rh159 and UL148, pInducer10-miR-RUP-PheS (54) (a generous gift of Stephen J. Elledge, Harvard Medical School, Addgene #44011) was digested with NotI and MluI to remove the miR-30 cassette and re-assembled using oligo RFP_stitch in Gibson reaction (55) to yield pIND-RFP. Plasmid pTRE3G-dTomato was assembled by Vector Builder. The TRE3G promoter was PCR-amplified with primers TRE3Gvb_Fw and TRE3Gvb_Rv and assembled into EcoRV-digested pSP72 by Gibson reaction to yield pSP72-TRE3G, which was sequence-verified using universal primer SP6. Following the example of Stinski and coworkers (56), the *crs* of the minimal CMV promoter within TRE3G was mutated from C<u>GTT</u>TAGTGAACCGT to C<u>AGG</u>TAGTGAACCGT by overlap extension PCR using primers TRE3G_crsmut_Fw and TRE3G_crsmut_Rv (57). Finally, the Δ*crs* TRE3G promoter was digested out of pSP72-TRE3G using NheI/AgeI and ligated into NheI/AgeI-digested pIND-RFP to yield pOUPc-RFP. pOUPc-UL148HA was constructed by PCR amplifying the UL148HA CDS from plasmid pcDNA3.1-UL148HA using primers UL148HAgibs_Fw and HAgibs_Rv and Gibson-assembling the product into AgeI/MluI-digested pOUPc-RFP. pOUPc-Rh159HA was constructed by PCR amplifying the Rh159HA CDS from plasmid pcDNA3.1-Rh159HA using primers Rh159HAgibs_Fw and HAeco_gibs_Rv and Gibson-assembling the product into AgeI/MluI-digested pOUPc-RFP. pOUPc-UL148HA and -Rh159HA were sequence-confirmed using primers CMVcrsnull_Fw and Ubc_Rv.

#### Lentivirus vector transduction

To generate stable i148^HA^ and i159^HA^ cell populations, replication defective HIV-1 based lentivirus vector particles were generated from pOUPc-UL148HA or -Rh159HA, as described previously (52). Briefly, 5 × 10^5^ 293T cells per well of a six-well cluster plate were co-transfected with pOUPc-UL148HA or pOUPc - Rh159HA, together with psPAX2 (Addgene #12260) and pMD2.G (Addgene #12259) using *Trans*IT-293 reagent (Mirus Bio, Inc.) as per the manufacturer’s instructions. Supernatants collected at 2 and 3 d post-transfection were combined, filtered through a 0.45 µm cellulose acetate syringe filter (Corning, Inc.), added to complete DMEM growth medium supplemented with 8 μg/mL polybrene (Sigma Aldrich) and applied to subconfluent ARPE-19 monolayers. The next day, medium was removed and the cells were washed three times with Dulbecco’s PBS (PBS: 2.7 mM KCl, 1.5 mM KH_2_PO_4_, 137 mM NaCl, 8.1 mM Na_2_HPO_4_, pH 7.4). Starting at 2 d post-transduction, cells were serially-passaged in medium containing 2 μg/mL puromycin HCl until resistant cells grew out.

#### Metabolic labeling

i159^HA^ or i148^HA^ cells were seeded at 2 × 10^5^ cells per well in a 24-well cluster plate in Gibco OptiMEM reduced serum medium (Thermo Fisher) supplemented with 2.5% tetracycline (tet)-free FBS (Clontech #631101). The following day, medium was replaced with 2.5% tet-free FBS OptiMEM supplemented with either 100 ng/mL doxycycline hyclate (dox, Sigma Aldrich #D9891, added from a 1000*×* stock) or the addition of 0.1% (vol/vol) of sterile water to control for volume of dox stock solution (mock induction). At 24 h post-induction, cells were washed twice in PBS supplemented with 1 mM CaCl_2_ and 0.5 mM MgCl_2_ and then incubated for 1 h in starving medium (DMEM lacking methionie, cysteine, and glutamine, (Gibco #21013024, supplemented with 5% dialyzed FBS, Sigma #F0392, and 2 mM glutamine). Cells were then pulse-labeled in starving medium containing 150 µCi/mL ^35^S-Met/Cys (PerkinElmer #NEG772) for 30 min. Dox or mock treatment was maintained throughout the starving and pulsing steps. As a positive control for translation shutdown, i159^HA^ ARPE19 cells were treated with 2 µM thapsigargin (Sigma #T9033) or 0.1% dimethyl-sulfoxide (DMSO) as a carrier-control at 1 h prior to Met/Cys starvation, and treatment was maintained throughout the starvation and pulse-labeling steps. Following pulse-labeling, cells were washed three times in PBS containing 1 mM CaCl_2_ and 0.5 mM MgCl_2_, and then immediately lysed in *2 ×* Laemmli buffer [120 mM Tris pH 6.8, 4% SDS, 20% glycerol, 0.02% bromophenol blue]. Beta-mercaptoethanol was then added to a final concentration of 5% vol/vol, and the samples were heated at 95˚C for 10 min. Equal volumes of lysate were resolved by sodium dodecyl sulfate polyacrylamide gel electrophoresis (SDS-PAGE) on 12% acrylamide NuPAGE Bis-Tris precast gels (Invitrogen #NP0321) according to manufacturer’s instructions. Gels were dried and exposed to a phosphor screen for 24 h before results were captured using an Amersham Typhoon IP scanner (GE Heathcare). Relative signal per lane was calculated using Bio-Rad 1-D analysis software by reading the signal volume (counts*mm^2^) in each lane. Lane signals were normalized either to (i) the DMSO treatment condition or (ii) each respective non-treatment condition.

### Cell viability assay

1 × 10^5^ i148^HA^ or i159^HA^ ARPE-19 cells per well were seeded in a 24 well cluster plate and incubated overnight. Medium was exchanged for complete DMEM containing 100 ng/mL doxycycline and incubated for 24 h. Cells were then trypsinized, transferred to 1.5 mL microfuge tubes and spun down at 400 × *g* for 5 min. Cell pellets were resuspended in 100 µL fresh medium and combined with 100 µL of PBS containing 0.4% Trypan Blue (Bio-Rad), mixed thoroughly and counted for viability (trypan blue exclusion) and total cell number using a hematocytometer (Bright-Line).

### Doxycycline induction of UL148 and Rh159 from stably transduced ARPE-19 cells

For each well of a 24 well cluster plate, 2 × 10^5^ cells of iUL148 or iRh159 ARPE19 cells were seeded in 500 µl OptiMEM medium containing 2.5% tetracycline(tet)-free FBS (Clontech #631101). Following a 24 h incubation at 37˚C, medium was replaced with fresh 2.5% tet-free FBS/OptiMEM supplemented with 100 ng/mL doxycycline hyclate (dox, Sigma Aldrich #9891). Where indicated, parallel wells of ARPE-19 cells were incubated for 4 h in the presence of 200 nM thapsigargin (Tg) prior to harvest. At the indicated times post treatment, cells were washed in PBS and lysed for 1 h at 4˚C using 50 µl per well of lysis buffer (1% Triton X-100, 400 mM NaCl, 0.5% sodium deoxycholate, 50 mM HEPES pH 7.5) supplemented with 1 *×* protease inhibitor cocktail (Cell Signaling Technology). Lysates were collected and spun down at 18,000 × *g* for 30 min at 4˚C. Protein concentrations of supernatants were measured using the BCA assay (Thermo Pierce), normalized, and subjected to western blot.

### siRNA treatments

2 × 10^5^ HFFTs per well of a 24-well plate were reverse transfected with 5 pmol per well of a Dharmacon siGENOME SMARTpool specific for human PERK (EIF2AK3, M004883-03-005) or with a non-targeting control SMARTpool (D-001206-14-05), using 4.5 µL of Lipofectamine RNAiMAX per well, as described previously (16). Briefly, siRNA transfection complexes in OptiMEM medium were added to wells prior to applying freshly trypsinized HFFT suspended in 0.45 mL of DMEM containing 8% FBS, 25 μg/mL gentamicin, and 10 μg/mL ciprofloxacin-HCl. 24 h post-seeding, cells were infected with the indicated viruses at an MOI of 1 TCID_50_ per cell. Sequences of the siRNAs in each SMARTpool are provided in Table 2.

### Xbp-1 splicing assay

HEK-293T cells were seeded into 24-well plates for overnight culture and transfected once they reached 80-90% confluency using the TransIT 2020 reagent (Mirus, Inc.), with each well receiving 1 µg plasmid DNA carried by 3 µL of the transfection reagent. 48 h post transfection, cells were harvested and total RNA was extracted using the Qiagen RNeasy mini kit as per the manufacturer’s protocol, including the optional column DNAase digestion step. cDNA was generated from 1 µg RNA using the qScript™ cDNA Synthesis Kit (Quantabio, Cat# 95047-100) in a 20 µL final reaction volume. One microliter of the resulting cDNA solution was then used as template for a PCR reaction using primers Xbp-1_FWD and Xbp-1_REV (Table 1). In the context of infection (Fig. 4), detection of *IE2* (*UL122*) mRNA was included as indicator of HCMV infection. The *IE2* primer pair was designed using PrimerQuest software (Integrated DNA Technologies, Coralville, IA), and includes one oligonucleotide whose priming site spans the junction of exons 3 and 5 (Table 1).

### Quantitative reverse-transcriptase PCR

mRNA levels were quantified using reverse-transcriptase qPCR (RT-qPCR). For these experiments, 2 × 10^6^ HFFT cells per well were seeded in a 6 well cluster plate, incubated overnight and subsequently infected at MOI 1. At 24 hpi, inocula were removed and replaced with fresh medium. At 72 hpi, total RNA was extracted using a Qiagen RNeasy Mini Kit (Qiagen, Inc.), including the optional on-column DNase digestion step, as per the manufacturer’s instructions. cDNA was generated from 1 µg RNA using the qScript™ cDNA synthesis kit. For each qPCR reaction, 1 µl of cDNA was used as template in a 15 µL final reaction volume using iQ SYBR Green Supermix (Bio-Rad, Inc.) on a CFX96 Real Time PCR system (Bio-Rad). mRNA levels for each gene were measured in triplicate technical replicates per biological replicate, with a total of three independent biological replicates, and the 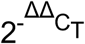 method (58) was used to determine quantitative estimates of relative gene expression, with all readings being normalized to *GAPDH* transcript levels. Canonical UPR responsive target genes were detected using previously validated qPCR primer pairs (29, 59), while levels of the viral *IE1 (UL123)* mRNA were also measured as an indicator of HCMV infection (Table 1). PCR efficiencies for primer pairs ranged from 91.1% - 99.0% for the indicated UPR target genes and were 93.4% for *GAPDH* and 90.7% for *IE1*. Statistical Analyses were done using GraphPad Prism software, version 6.0h (GraphPad, Inc., La Jolla, CA).

### Antibodies used in this study

The following rabbit monoclonal antibodies (mAbs) from Cell Signaling Technology, (Danvers, MA) were used: ATF4 clone D4B8 (cat. #11815S), PERK clone C33E10 (cat. #3192S), Phospho-eIF2α Ser51 clone D9G8 (cat. #3398S), total eIF2α clone D7D3 (cat. #5324S). IE1 was detected using mouse mAb clone1B12 (Gift of Thomas Shenk, Princeton University), beta-actin using a rabbit mAb, (Li-Cor Biosciences, Inc.; cat. #926-42210), a rabbit polyclonal anti-HA epitope antibody (Bethyl Laboratories, Inc., Montgomery, TX), and a previously described rabbit polyclonal serum specific for UL148 (19).

### Western blotting

For detection of all proteins other than phospho-Ser51 eIF2α (see below), western blotting was carried out as previously described (16, 19, 52). Briefly, cells were lysed at 4˚C for 1 h in lysis buffer [1% Triton X-100, 400 mM NaCl, 0.5% sodium deoxycholate, 50 mM 4- (2-hydroxyethyl)-1-piperazineethanesulfonic acid (HEPES) pH 7.5 supplemented with 1 × protease inhibitor cocktail (Cell Signaling Technology)]. Lysates were clarified by centrifugation at 18,000 *× g* for 30 min at 4˚C, combined with an equal volume of 2 × Laemmli buffer containing 10% betamercaptoethanol, heated at 85˚C for 10 min prior to being resolved by SDS-PAGE on 10% acrylamide gels and transferred to nitrocellulose membranes (Whatman Protran^®^, 0.45 µm pore size). Efficient transfer was confirmed using Ponceau S staining (not shown). All subsequent blocking, washes, and incubation steps were performed with gentle rocking. Membranes were blocked using a solution of 5% powdered milk (PM) in PBS containing 0.01% Tween-20 (PBST), (PM-PBST). Unless otherwise noted, all antibodies were applied to membranes in PM-PBST, as a 1:1000 dilution, or for IE1 mAb, a 1:200 dilution of the hybridoma supernatant, and incubated overnight 4˚C, or for 1 h at room temperature. Following three 5 min washes in 1 × PBS, IRDye-800 conjugated donkey anti-rabbit or anti-mouse secondary antibodies (Li-Cor, Inc.) were applied at 1:10,000 in PM-PBST and incubated for 1 h. After 3 washes in PBST, immunoreactive polypeptides were detected, and where applicable, quantified using a Li-Cor Odyssey Imager (Li-Cor Biosciences). For detection of Ser51 phosphorylated eIF2α, a protocol from the laboratory of David Ron (Cambridge Institute for Medical Research, United Kingdom) was used. The differences from our standard procedures were as follows: Membranes were blocked for 2 h at room temperature in a solution of 5% bovine serum albumin (BSA) in PBS containing 0.01% Tween-20, followed by a second 10 min blocking step in PM-PBST. Following three washes in PBS, membranes were incubated overnight in a solution of phospho-eIF2α antibody diluted 1:1000 in PBS supplemented with 5% BSA.

## ACKNOWLEDGMENTS

This project was supported by NIH Grants R01-AI116851 and P30GM110703. Its contents are solely the responsibility of the authors and do not necessarily represent the official views of the NIAID or the NIGMS. C.C.N. was supported by a Malcolm Feist Predoctoral Fellowship from the Center of Cardiovascular Diseases and Sciences at LSU Health Sciences Center, Shreveport. We are grateful to Thomas Shenk (Princeton University, Princeton, NJ, USA), Christian Sinzger (University of Ulm Medical Center, Ulm, Germany), and Klaus Früh (Oregon Health Sciences University, Beaverton, OR, USA) for generously providing reagents.

